# Multiple Monkey Pose Estimation Using OpenPose

**DOI:** 10.1101/2021.01.28.428726

**Authors:** Salvador Blanco Negrete, Rollyn Labuguen, Jumpei Matsumoto, Yasuhiro Go, Ken-ichi Inoue, Tomohiro Shibata

**Affiliations:** Department of Human Intelligence Systems, Graduate School of Life Science and Systems Engineering, Kyushu Institute of Technology, Kitakyushu City, Fukuoka, Japan; Systems Emotional Science, University of Toyama, Toyama, Japan; Cognitive Genomics Research Group, Exploratory Research Center on Life and Living Systems (ExCELLS) National Institutes of Natural Sciences, Okazaki, Aichi, Japan; Department of System Neuroscience, National Institute for Physiological Sciences Okazaki, Aichi, Japan; Systems Neuroscience Section, Department of Neuroscience, Primate Research Institute, Kyoto University, Inuyama, Aichi, Japan

**Keywords:** OpenPose, Monkey, PAF

## Abstract

This paper proposes a system for pose estimation on monkeys, in diverse and challenging scenarios, and under complex social interactions by using OpenPose. In comparison to most animals used for research, Monkeys present additional difficulties for pose estimation. Multiple degrees of freedom, unique complex postures, intricated social interactions, among others. Our monkey OpenPose trained model is robust against these difficulties. It achieves similar performance as in human pose estimation models, and it can run in Realtime.

## I. INTRODUCTION

Scientists use animal models for studies that include human deceases, genetic models, medical trials, and behavioral studies. Animals are even used by the industry to test products' safety, such as foods and cosmetics. Among the different animal models, nonhuman primates more closely mirror human physiological and behavioral features.

A popular approach for monkeys' behavior analysis is to categorize the monkeys' activities and behaviors manually. Nevertheless, this approach requires trained personal familiar with monkey physiology. This approach is too costly, and results can change from evaluator to evaluator, making evaluation imprecise and difficult to reproduce. For these reasons, automatic pose estimation is of great importance. Animal pose encodes information regarding behavior. Moreover, the posture can be measured quantitatively.

Commercial marker-based pose estimation systems readily exist [1]. However, physical markers are not desirable since they can intrude and perturb animals' behavior, especially monkeys. Monkeys tend to remove the markers; monkey's hair causes occlusion and makes markers difficult to attach. Interaction with the environment and other monkeys may also detach the markers. Moreover, some of these systems rely on expensive equipment, and its use is often restricted to laboratory caged setups.

This work presents a multiple monkey markerless pose estimation system that uses OpenPose, originaly developed for human pose estimation [2]. Monkey pose estimation can be used in various environments, and there is no restriction on the number of subjects.

## II. RELATED WORK

A markerless approach to monkey pose estimation has been realized through the use of multiple cameras with deep sensing capabilities, where a skeleton model is fitted into a captured monkey 3D image [3]. But this approach requires calibration of the cameras, the number of subjects is restricted, and it requires human intervention for correction when the system fails. Similarly, [4] requires numerous cameras for 3D reconstruction; the number of subjects is limited and restricted to a laboratory environment.

It has been shown that a deep learning approach is feasible [5][6][7]. Although monkey pose could be estimated accurately, these works are limited to a single subject. Thus the analysis of social interactions was not possible. In a recent work [7], DeepLabCut (DLC) was used for monkey pose estimation, DLC is a DeepLearning framework for body feature detection on single animals, although a multi-animal option will be available in the future [8]. The mulitple subject feature is currently in development only available as a beta, so a comparison was not possible. By using transfer learning from a human model, chimpanzee dense pose estimation is possible [9]. However, at least for evaluation manual labeling is necessary. Dense pose annotation is very challenging and a 3D artist-created reference model is necessary for labeling. The authors even mention the omission of dense annotations for body parts due to its complexity for annotators. Part affinity fields have been used on honeybees and mice pose estimation and tracking [10]. However, the experiments is done in a highly controlled laboratory environment; furthermore, the skeletons and behaviors of honeybees and mice are simple in comparison to monkeys.

This work takes advantage of similarities in skeleton structure between humans and monkeys. OpenPose originaly used for humans is employed. OpenPose uses Part Affinity Fields (PAF); unlike feature detection based on location, the PAF encodes the limb orientation [2].

## III. METHOD

### A. Dataset

The MacaquePose monkey dataset was used. A total of 13,083 RGB images containing a total of 16,393 monkeys were used to train the model [7]. The monkeys exhibit a variety of poses and social interactions. They feed, groom, sleep, fight, climb, swim, etc. Multiple backgrounds, occlusion, noise, and other artifacts are present, which makes the dataset challenging. From all the images, 93.75% are used for training and 6.25% for validation. Table 1 gives more information regarding the distribution of the dataset.

**Table 1.** Training and validation set distribution

|  | Train | Val | Total |
| --- | --- | --- | --- |
| Images | 12265 | 818 | 13083 |
| Images % | 93.75 % | 6.25 % |  |
| Monkeys | 15373 | 1020 | 16393 |
| Monkeys % | 93.77 % | 6.22 % |  |

Each monkey has 17 labels for each body feature as described in Figure 1. If the body feature is present, it is described by its (x,y) coordinates; and an additional bit indicates whether the body feature is visible or occluded. The neck body feature is not labeled but is calculated in relation to the other body features.

**Figure 1.**
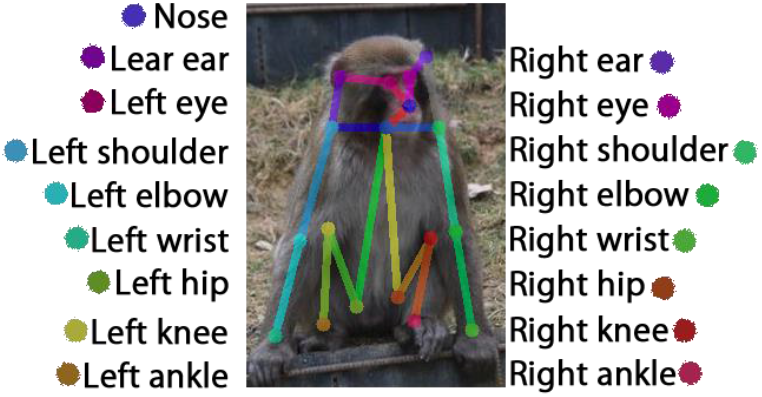
Sample data anotated 17 body features.

### B. Network Architecture

We use hyperpose an open-source implementation part of the TensorLayer project [11]. TensorLayer is part of Google's TensorFlow Framework for machine learning and deep learning. In this work, ResNet18 is used as a backbone for the network [11]. Although the use of a deeper backbone like ResNet50 could yield better results, ResNet18 allows the network to be lightweight. It reduces the training, execution time and computational requirements for it to operate, making it more accessible. Following the same line of thinking, the head of the network, is a lightweight implementation of OpenPose [12]. The network takes body feature related labels and mask for each monkey. As an output, the network generates score maps; then, it estimates preliminary part affinity fields. After a refinement stage the skeleton of the monkeys is generated.

Once the model is trained, it is exported to the Open Neural Network Exchange (ONNX) format that allows it to run in the Nvidia C++: TensorRT high-performance deep learning inference framework.

### C. Evaluation and Training

Average Precision (AP) is the metric used to evaluate the model. Standard augmentation methods like random rotation, shifts, and flips were used during training. A graph of the training Loss is also provided. The network takes images with a maximum dimension of 640 pixels, either on the height or width. The model is trained on a Nvidia GeForce GTX TITAN X graphics card for up to 100,000 iterations and a batch size of 8. The required time to train the network is approximately 24 hours. The same computer set up is use for evaluation and inference.

## IV. RESULTS

The model was trained up to 100,000 iterations taking around 24 hours. As Figure 2 shows, the Loss significantly decreased during the first 10,000 iterations. Afterward, the Loss keeps dropping at a slower but steady rate. The graph also shows a comparison against the same network trained with the human MSCOCO 2017 dataset [13]. The graph shows the monkey model training performance is comparable to the human one; indeed, the monkey model performs slightly better. It is remarkable to see the monkey model achieving close results to the human model since the monkey dataset is approximately ten times smaller. The COCO training set contains over 100,000 person labeled instances. The better results on the monkey model could be explained by the MSCOCO dataset being more challenging. The MSCOCO dataset, in addition to humans, contains numerous objects and animals, while the monkey dataset, for the most part, only contains monkeys.

**Figure 2.**
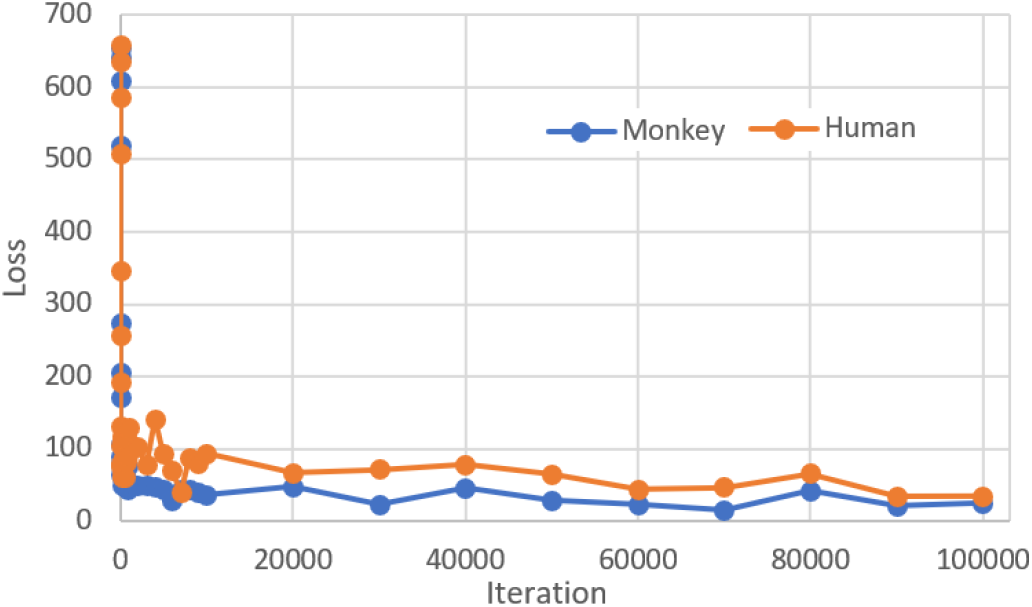
Monkey and Human Loss for training

Figure 3 shows the results during the training of the PAF and score maps after 100,000 iterations. Figure 3 (c) shows that the model can generate the PAF even in challenging situations and with multiple subjects. The generated PAF closely reassembles the ground truth PAF (b). When comparing against the ground truth, it is important to consider that the ground truth does not reflect the ease of visibility or occlusion. In the example, it can observe that extremities and occluded body features have lower confidence for PAF and score maps. It can also be observed that facial features and easily visible features have a higher confidence score. These results are in accordance with previous works for single monkey subjects [6].

**Figure 3.**
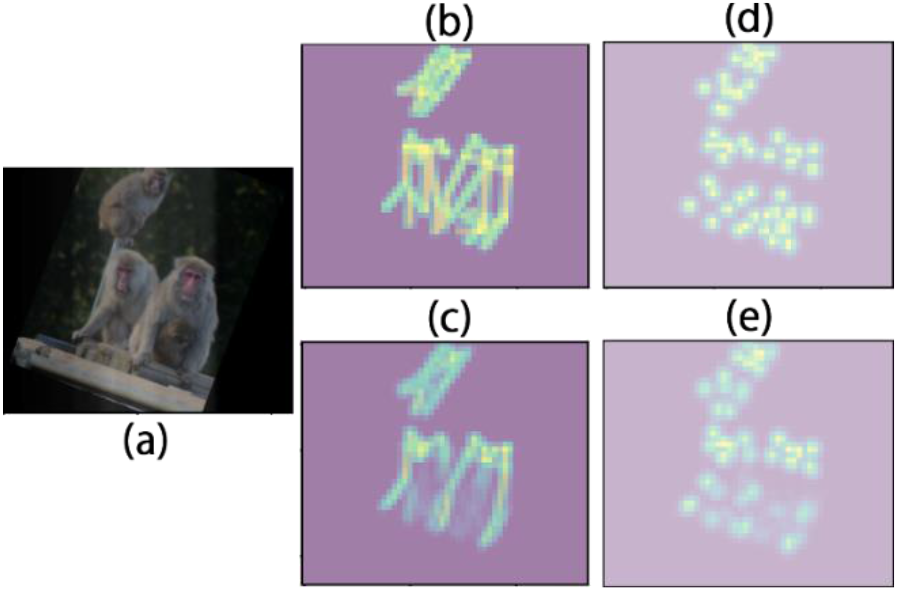
Results after 100000 iterations a) input image with augmentation (b) PAF ground truth (c) PAF result (d) score maps ground truth (e) score maps result

### A. Evaluation

To evaluate the performance of the trained model, AP was used as a metric. The monkey trained model is also compared against the same network trained on the MSCOCO2017 human dataset. The results reported on the original OpenPose are also included as a reference [2]. The OriginalOpen pose implementation and backbone is different, so a fair comparison is not possible. The original OpenPose backbone is based on the VGG-19 while our monkey trained model is based on ResNet18 and uses a lightweight implementation of OpenPose. Additionally, the original OpenPose is evaluated on the MSCOCO2016 dataset.

Table 2 shows the result of Ap^50^, Ap^75^, Ap^M^, Ap^L^. Twenty-nine randomly selected images from the evaluation set were selected to evaluate the Monkey model. Similarly, to evaluate the human pose trained model, eleven randomly selected images from the evaluation set were selected. Compared to the trained network on humans, the monkey network achieves better performance in the four Ap; this can be attributed to the more challenging human dataset. Compared to our monkey model, the original OpenPose has better performance.

**Table 2.** Evaluation Ap

| Network | Dataset | $Ap^{50}$ | $Ap^{75}$ | $Ap^M$ | $Ap^L$ |
| --- | --- | --- | --- | --- | --- |
| Lightweight Openpose + ResNet18 | Monkey Dataset | 76.2 | 38.0 | 35.5 | 43.2 |
| Lightweight Openpose + ResNet18 | MSCOCO 2017 | 58.7 | 10.4 | 18.2 | 28.1 |
| Original OpenPose | MSCOCO 2016 | 83.4 | 66.4 | 55.1 | 68.1 |

### B. Visual assessment

Images from the evaluation set were also analyzed Figure 4, 5 and 6. These images were not seen during training. Figure 4, in addition to the final result, also shows the scoremaps and PAF. The output score maps confirm that the network can infer in unseen images monkey body features and its relation, represented by the generated PAF.

**Figure 4.**
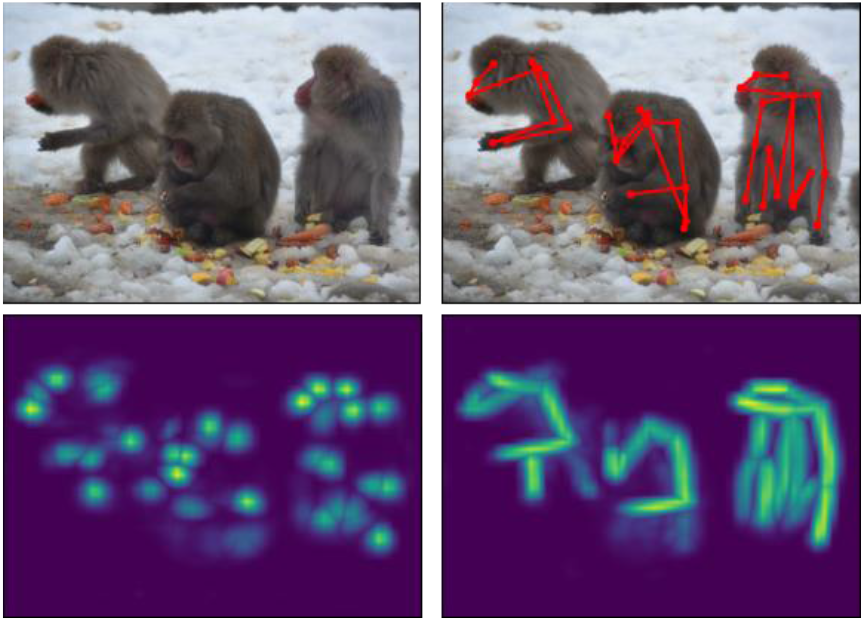
Input image (top left) score map prediction (bottom left) PAF prediction (bottom right) final output (top right)

**Figure 5.**
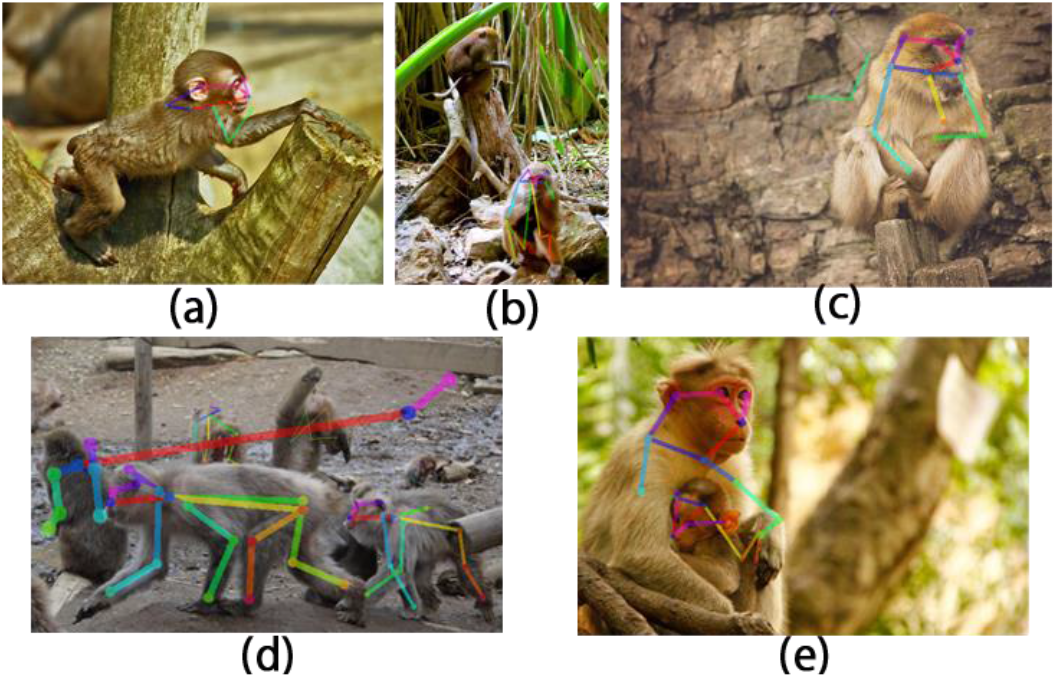
Sample failure cases: (a) missing body parts (b) missing monkey detection (c) false detection on the background (d) random detections (e) same body part detection shared by multiple individuals and association of bodyparts form different individuals to the same skeleton.

**Figure 6.**
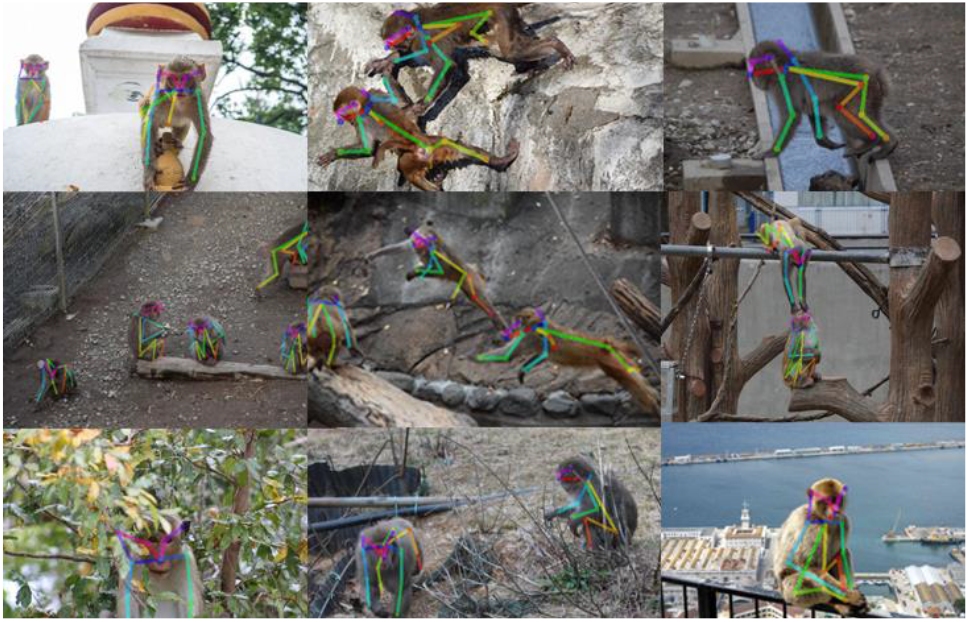
Success detection sample: challenging and varied environments, multiple individuals, variety of activities, occlusion, social interactions.

#### 1) Failure cases

Some of the most comon falilure cases are ilustrated in Figure 5, a) although a monkey is detected, some of its body parts are not; in most cases, this could be attributed to a low confidence score on the missing body features. b) A monkey is not detected; this tends to happen more when facial body features are not visible. In some of these cases, monkeys are tricky to spot even for a human due to bending with the environment or small scale. c) In a few cases, some body parts of a nonexisting monkey are detected in the background. d) False body features of an existing monkey are detected on the background; in most instances, the false detected body feature is close to the subject, but in a few cases, it could look like a random detection. e) A skeleton prediction takes body parts from different monkeys, or in some cases, two generated monkey skeletons share a single body feature. This errors tends to occur when monkeys are closely interacting.

#### 2) Success cases

Figure 6 shows a sample of successful detections on unseen images from the dev set. The sampled images display various activities that include eating, playing, jumping, crawling, and standing. The backgrounds are rich and varied, ranging from natural sceneries with plants and trees to a city with a sea view.

### C. Real time performance

The trained model was exported to the Open Neural Network Exchange (ONNX) format, and ran in the Nvidia C++: TensorRT high-performance deep learning inference framework. Our trained model in the ONNX format size is of 31 MB. In our testing we were able to run the model in real time using the webcam stream. The hyperpose open source project reports being able to run at 60 fps using the default OpenPose with the ResNet18 as a backbone and a resolution of 432 x 368 images. Our trained model uses a lightweight version of OpenPose so it should yield a superior performance.

## V. CONCLUSION

In this work, we have trained an OpenPose neural network using the MacaquePose monkey dataset. The trained model is capable of detecting monkey body features and generating PAF on unseen images. The final output is the monkey's posture; there are no restrictions on the number of monkey subjects in the image. It is robust against colussion and social interaction, and it works on challening backgrounds.

The model was exported to the ONNX format allowing the model to run in real-time. Better results could be achieved on videos by using information from contiguous frames, or by tracking the subjects. Moreover, the model could be improved by using a deeper network and transfer learning.

The proposed model was shown to achieve comparable performance to a model trained on a human dataset. The pose estimation achieved could be used to study the monkeys' behavior. The multiple subjects' pose estimation capabilities open the possibility to study complex monkey social interactions.

## Supporting information

Pose detection group of monkeys

## ACKNOWLEDGEMENTS

This work was supported by the Cooperative Research Program of Primate Research Institute, Kyoto University, the Grant-in-Aid for Scientific Research from Japan Society for the Promotion of Science (JSPS Grant Numbers JP16H06534), the grant of Joint Research by the 228 National Institutes of Natural Sciences (NINS) (NINS Program No. 01111901), and the Mexico National Council of Science and Technology, CONACYT.

